# Endogenous suspension and reset of consciousness: 7T fMRI brain mapping of the extended cessation meditative endpoint

**DOI:** 10.1101/2025.09.06.674021

**Authors:** Winson F.Z. Yang, Akila Kadambi, Kilian Abellaneda-Pérez, Grace Mackin, Isidora Beslic, Ruby Potash, Terje Sparby, Matthew D. Sacchet

**Affiliations:** Meditation Research Program, Department of Psychiatry, Massachusetts General Hospital, Harvard Medical School, Boston, USA; Athinoula A. Martinos Center for Biomedical Imaging, Department of Radiology, Massachusetts General Hospital, Harvard Medical School, USA; Dept. of Psychiatry and Biobehavioral Sciences, David Geffen School of Medicine, University of California, Los Angeles, USA; Brain and Creativity Institute, Dornsife College of Letters, Arts, and Sciences, University of Southern California, Los Angeles, USA; Institut Guttmann, Institut Universitari de Neurorehabilitació adscrit a la Universitat Autònoma de Barcelona, Badalona, Spain; Department of Medicine, Faculty of Medicine and Health Sciences, Institute of Neurosciences, University of Barcelona, Barcelona, Spain; Northeastern University, Boston, USA; Steiner University College, 0260 Oslo, Norway; Department of Psychology and Psychotherapy, Witten/Herdecke University, 58455 Witten, Germany; Integrated Curriculum for Anthroposophic Psychology, Witten/Herdecke University, 58455 Witten, Germany

**Keywords:** advanced meditation, extended cessation, consciousness, subcortical, brainstem, functional connectivity, chemoarchitecture

## Abstract

Extended cessation (EC), an advanced meditative state in which consciousness is volitionally suspended and later reset with immense mental clarity, equanimity, and peace, offers an endogenous model for investigating the mechanisms of consciousness. Using ultra-high-resolution 7T fMRI with dense within-subject sampling (N=3), we quantified whole-brain activity, functional and effective connectivity, cortical gradients, and eigenmodes, and related them to chemoarchitecture and cognitive maps. EC is marked by increased activity in unimodal regions, down-regulation in transmodal regions, subcortex, and brainstem, an expansion of the principal gradient, and decrease in low-order global eigenmodes. Cognitive decoding linked EC to heightened perceptual clarity and attention, least with mental suffering, and co-varied with histaminergic H receptors topology. These findings challenge predictions of Global Neuronal Workspace and Integrated Information Theory, while supporting the Active Inference Framework. More broadly, EC demonstrates that consciousness can cease without global suppression, suggesting a potential “reset” mechanism that fosters equanimity and the potential for flourishing.

## Introduction

In most scientific and clinical frameworks, wakefulness is assumed to entail ongoing subjective consciousness^1^. Advanced meditative states challenge this assumption by showing that consciousness can be radically thinned to minimal ‘content-free’ awareness^2,3^ or even entirely suspended momentarily^4–7^ or for an extended duration^8^. These findings highlight the extraordinary plasticity of consciousness, raising fundamental questions for theories of consciousness, including Global Neuronal Workspace, Integrated Information Theory, and Active Inferences Framework, and their relevance to human flourishing^9,10^.

Advanced meditation situates these rare states within a broader scientific agenda: understanding consciousness as a trainable and reconfigurable capacity across meditative development^11,12^. This systematic mental training can lead to meditative endpoints, variously described as “awakening”^13,14^, “beatitude”^15^, “salvation”^16^, or “enlightenment”^13,14^, that entail profound shifts in perception, emotion, and self-processing. Studying such endpoints offers not only a natural experiment for testing theories of consciousness but also insights into how such they can induce a “reset” of consciousness that fosters equanimity, resilience, and well-being, all of which are critical priorities in global mental health^17–19^.

Here, we study extended cessation (EC), a rare meditative endpoint where practitioners report temporary suspensions of ordinary consciousness and mental activities^4,8^. EC can vary from minutes to days, is entered and exited volitionally, and is frequently followed by profound psychological aftereffects, including extreme sense of relief, clarity, openness, peace, equanimity, heightened sensory sensitivity, significant reduction in mental suffering, reduced repetitive thoughts, stoppage of negative self-talk, and an absence of inner narration^8,18,19^. Unlike non-conscious states induced by pharmacological agents (e.g., anesthesia or psychedelics)^20,21^, EC is unique in being endogenous, reversible, precisely timed, and followed by heightened well-being, making it an excellent natural experiment for probing the mechanisms of consciousness. One canonical form of EC found in Theravada Buddhism is Nirodha Samāpatti^8,22–24^. Here, we investigate the broader neurobiological principles of EC without making definitive claims about its doctrinal interpretation^4,6,7^. Please refer to Laukkonen et al. for a recent discussion of the historical and theoretical context of Nirodha Samāpatti^8^. It is noteworthy that Nirodha Samāpatti is considered to be among the highest, or the highest, meditative attainment in some interpretations of Theravada Buddhism^25,26^.

From a neuroscience perspective, studying EC offers a powerful opportunity to explore the fundamental nature and mechanisms of consciousness. A central limitation in current consciousness research is the absence of an endogenous and reversible method to suspend awareness while preserving physiological stability^20,21^ and enabling prospectively timed re-emergence^8,23^. EC addresses this gap^7,8^. Compared to sleep, anesthesia, and disorders of consciousness, EC constitutes a fourth experimental context in its uniqueness self-initiate and control consciousness. This enables precise examination of transition dynamics at entry and exit and the mechanisms that dismantle and reinstate conscious access. Additionally, the notable aftereffects of EC described above may relate to significant neuroplasticity and reorganization of the brain, potentially informing techniques to enhance cognitive function, emotional regulation, overall wellbeing and happiness, and thriving.

Current competing frameworks of consciousness make testable predictions for EC. Global Neuronal Workspace (GNW) predicts a collapse of long-range connectivity and reduced feedback connectivity during suspension of consciousness^27^ while Integrated Information Theory (IIT) predicts a decrease in network differentiation during non-conscious states^28^. In contrast, the Active Inference Framework (AIF) predicts down-weighting of sensory precision via thalamo-cortical gating during non-conscious states^29^. The distinct features of EC allow comparison among theories about how consciousness is represented and suspended in the human brain.

Here we present the first neuroscientific, empirical investigation of EC, using ultra-high-resolution 7T functional magnetic resonance imaging (fMRI) to investigate the whole-brain activity, connectivity, and causal dynamics of network organization during EC. The ultra-high spatial resolution of 7T imaging allows for precise measurement of small but critical brain structures that are central to consciousness and arousal regulation, such as the thalamus, brainstem, and subcortical nuclei. This methodology is particularly powerful for the study of consciousness, where transitions between conscious and non-conscious brain states may be mediated by subtle shifts in deep-brain activity. Moreover, we used a novel and systematic neurophenomenological approach to rigorously examine the subjective experience of EC, enabling a rare convergence of first-person insight and third-person measurement.

This study provides important initial evidence for the neuroscience of EC and related fundamental aspects of consciousness, with potential therapeutic implications for human flourishing. In this study we sought to answer three key questions: (1) What brain reconfiguration accompanies voluntary suspension of consciousness?; (2) What are the causal brain dynamics of EC?; and (3) Which neurochemical systems are implicated, as inferred from spatial coupling to normative receptor distributions?

Our findings indicate that consciousness can be suspended without global cortical suppression. During EC, unimodal systems (e.g., visual and dorsal-attention networks) remain selectively engaged while transmodal hubs and thalamo-striatal circuits are down-regulated; the principal cortical gradient polarizes rather than flattens; and network–neurochemical coupling aligns with histaminergic (H) precision control of sensory gating. Meta-analytic decoding demonstrated that EC is functionally aligned with perception and attention, while being least associated with mental suffering and distress. This neuronal signature is most consistent with AIF, constrains GNW theory, and challenges IIT. This study demonstrates that the human brain can volitionally dismantle and reassemble conscious processing, offers a radical model for testing theories of consciousness, and highlights the possibility that consciousness can ‘reset’ and orient toward equanimity and flourishing.

## Results

### EC Phenomenology

Detailed demographic of participants (N = 3) are provided in Methods. Participant endorsement of EC phenomenology is presented in **Fig. 1**. Here, we summarize the key phenomenology for the setup, entry, exit, and afterglow of EC.

**Fig. 1.**
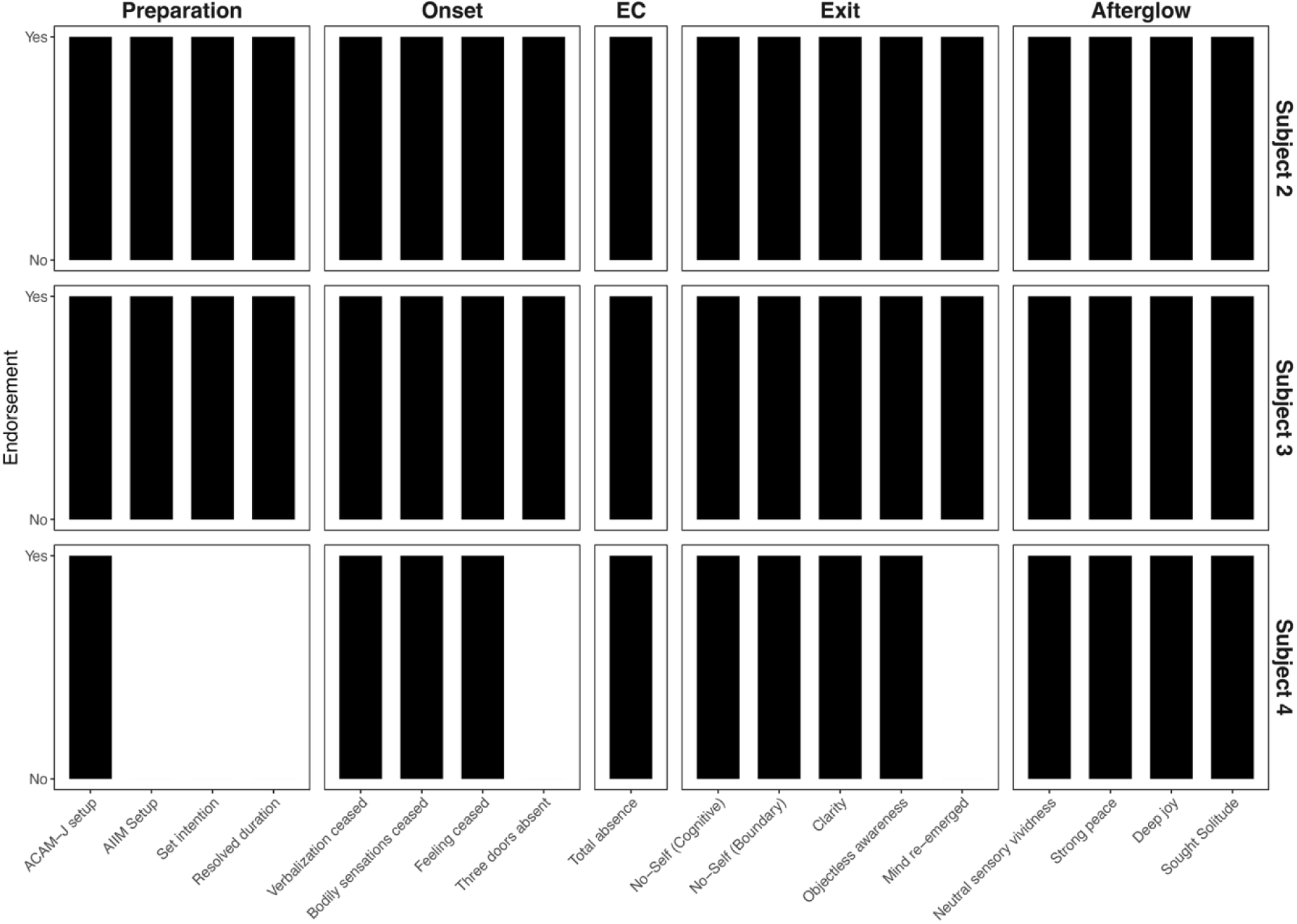
Endorsement of EC phenomenology for each participant. Participants all endorsed the phenomenological items of EC presented here, except Subject 4. Although previously trained to set up EC using ACAM-J, Subject 4 had since developed an alternative approach and no longer required ACAM-J for setup. As such, the phenomenology for various components throughout EC differ slightly for Subject 4. ACAM-J = Advanced concentration absorption meditation-jhana, AIIM = advanced investigative insight meditation

***Setup.*** Participants typically used advanced absorption concentration meditation (ACAM) to set up EC. ACAM is a series of meditative absorption states accompanied by blissful sensations, emotional happiness, and equanimity, followed by an increasing reduction of input from the physical environment and the reduction of contents of consciousness to a bare minimum experience. One participant who had previously trained using ACAM for setup, no longer necessarily needed ACAM to enter EC. Additionally, all but one participant incorporated elements of advanced investigative insight meditation (AIIM), that is, a neutral observation of phenomena, as part of their preparatory routine for EC.

***Onset.*** All participants reported that internal verbalization (thinking), body sensations, and feeling ceased during the onset of EC. With one exception, participants reported no primary experiential emphasis on either incongruence, transient nature of experience, or self-lessness (these are typically present during momentary cessations).

***EC.*** During EC, participants reported that typically all consciousness experience was completely offline.

***Exit.*** Upon emerging from EC, all participates reported that biographical memory and ego-centered cognitive processing were typically absent, and they did not perceive any separation between themselves and the environment. All participants reported a vivid clarity of mind and an experience of objectless awareness. All but one participant reported the return of mental activity in a specific sequence: first thought, then bodily sensation, and finally internal verbalization.

***Afterglow.*** After EC, all participants reported typically experiencing intense sensory vividness, accompanied by a strong and lasting peace. They all also reported typically being inclined to solitude, and many described a deep, abiding joy.

### Brain activity during EC: Regional homogeneity (ReHo)

Comprehensive results/summary tables of significant regions are included in the **Supplementary excel file**. Results are visualized in **Fig. 2**. Here, we only provide a summary of the results.

**Fig. 2.**
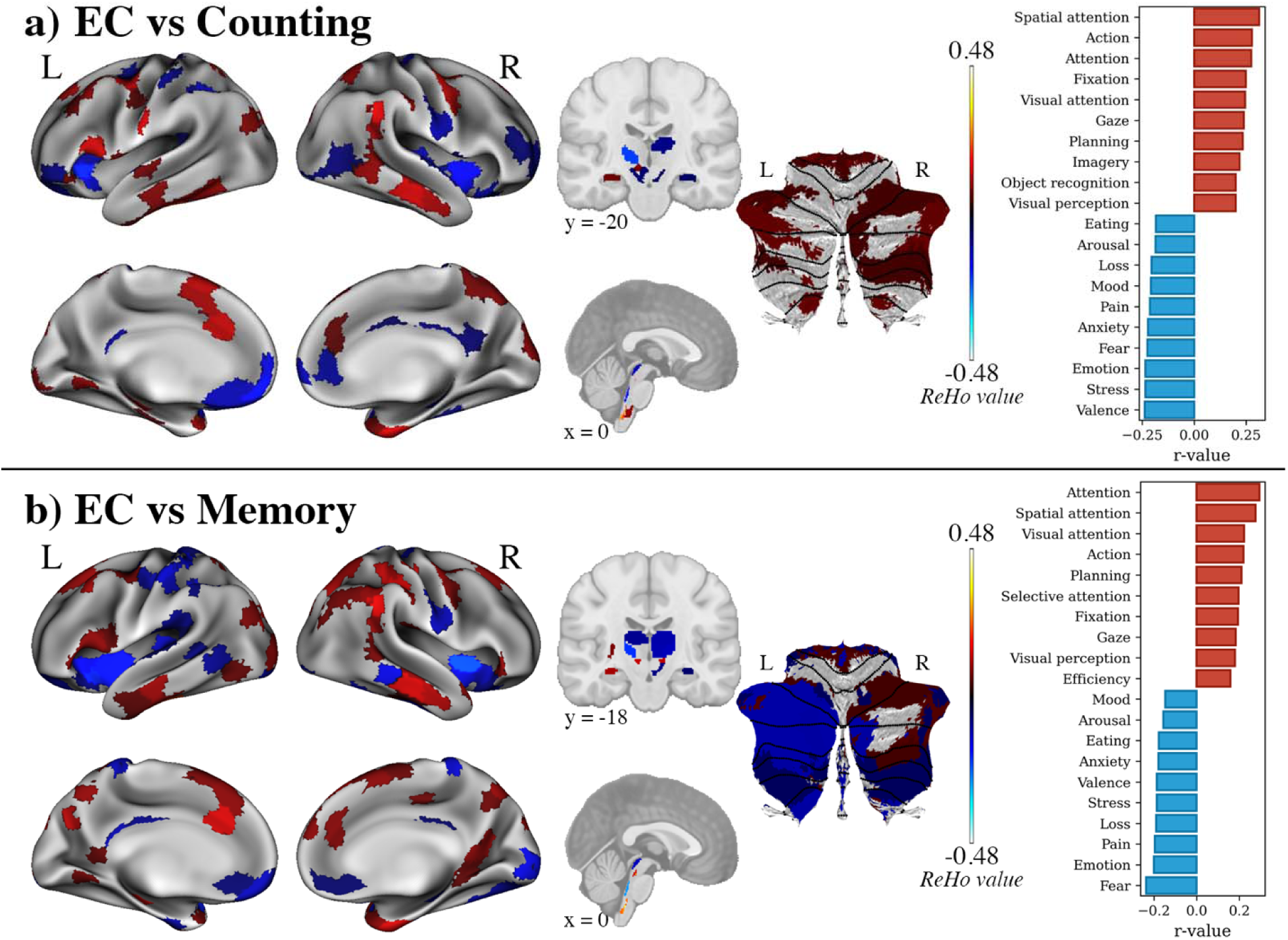
Differences in regional homogeneity values (ReHo) between EC and control conditions. **(a)** EC, compared to the counting control condition, demonstrated increased ReHo values in the bilateral visual, posterior, temporal regions, parts the brainstem, and cerebellum, and decreased ReHo in bilateral prefrontal regions and posterior cingulate cortex, largely associated with the default-mode network. Subcortical activity reveals decreased ReHo values across the caudate, thalamus, globus pallidus, amygdala, and brainstem. **(b)** Similarly, compared to the memory condition, EC displayed increased ReHo values in bilateral visual, posterior, and temporal regions, and decreased ReHo values in the prefrontal and insular regions, and posterior cingulate cortex. Subcortical regions primarily showed decreased ReHo values spanning the caudate, thalamus, globus pallidus, hippocampus, amygdala, parts of the brainstem, and cerebellum. Neurosynth decoding analysis revealed that these patterns of brain activity were related to cognitive processes associated with perception (e.g., attention, spatial attention, visual attention) and least likely with mental suffering and psychological distress (e.g., fear, stress, loss, pain).

***Across both counting and memory control conditions***. In the cortex, EC, compared to both counting and memory conditions, was associated with higher ReHo values in the bilateral visual and posterior cortices, frontal eye fields, temporal poles, left prefrontal, parietal, temporal cortices, and right dorsomedial prefrontal cortex; and lower ReHo values in the bilateral prefrontal, somatomotor, temporal, and orbitofrontal cortices, posterior frontal opercula and insula, left precuneus-posterior cingulate cortex, and right cingulate cortex.

In the subcortex, EC was primarily associated with decreased ReHo values. Regions include the ventoposterior (THA-VP) and right dorsoanterior (THA-DA) thalamus, left anterior putamen, right hippocampus, medial amygdala, anterior caudate, and globus pallidus. Increased ReHo was also seen in the left hippocampus, medial amygdala, and shell of the nucleus accumbens.

In the brainstem, EC was associated with higher ReHo values in the median raphe, raphe obscurus, left inferior medullary reticular formation (MRt), mesencephalic reticular formation, inferior olivary nucleus, viscero-sensory-motor nuclei complex, right alpha part of the parvicellular reticular nucleus, superior MRt, and superior olivary complex. EC was also associated with lower ReHo values in the dorsal raphe, left locus coeruleus, subcoeruleus, laterodorsal tegmental nucleus-central gray of the rhomboencephalon, right cuneiform nucleus, superior colliculus, and parabrachial pigmented nucleus complexes of the ventral tegmental area (VTA-PBP).

In the cerebellum, EC was associated with higher ReHo values in cerebellum regions 6 (divided attention) and 10 (autobiographical recall and interference resolution).

Additional significant regions of interest (ROIs) for each control condition are presented below.

***EC vs counting control condition.*** Additionally, EC, compared to the counting control condition, was associated with lower ReHo values in the left dorsoposterior thalamus (THA-DP), raphe magnus, left substantia nigra, and red nucleus.

***EC vs memory control condition.*** In the subcortex, EC compared to the memory control condition, was associated with higher ReHo values in the left posterior globus pallidus and putamen, and right nucleus accumbens. Decreased ReHo values were found in the bilateral lateral amygdala and right ventroanterior (THA-VA) thalamus and THA-VP. In the brainstem, higher ReHo values were found in the left isthmic reticular formation, pedunculotegmental nucleus, superior MRt, and right medial parabrachial nucleus; and lower ReHo values in the right inferior olivary nucleus and microcellular tegmental nucleus – prabigeminal nucleus. In the cerebellum, EC, compared to the memory control condition, was associated with lower ReHo in the cerebellar region 4 (associated with action observation).

Full correlations for the 123 cognitive terms are reported in the **Supplementary excel file.**

### Functional connectivity: Network-based statistics (NBS)

Comprehensive results/summary tables of significant regions are included in the **Supplementary excel file** and results are visualized in **Fig. 3a, b**. Here, we only provide a summary of the results.

**Fig. 3.**
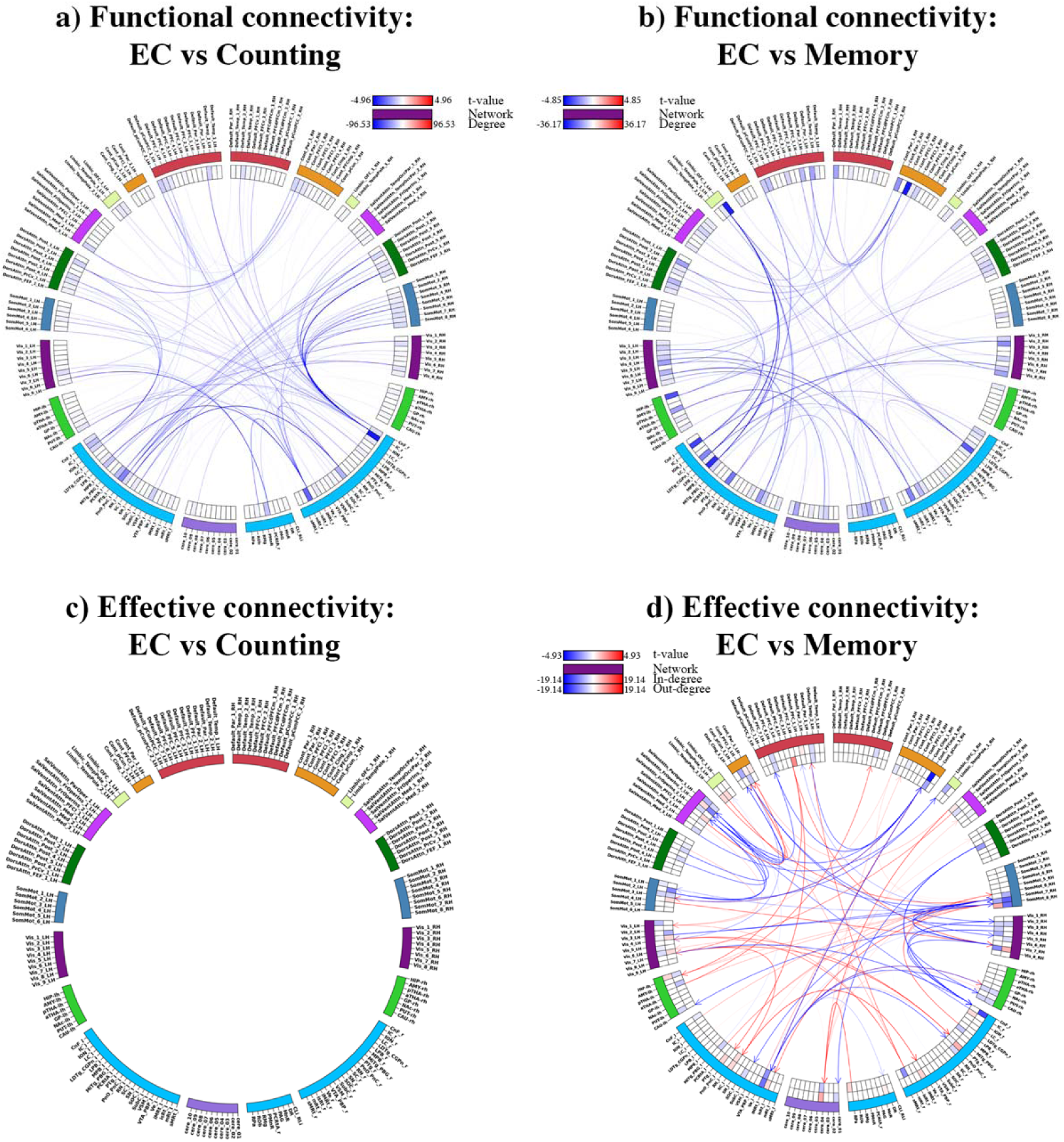
Differences in functional connectivity and effective connectivity between EC and control conditions. The figure illustrates **(a, b)** functional and **(c, d)** effective connectivity differences between EC and control conditions. Nodes represent brain regions, arranged based on their hemispheric locations and colored according to their network affiliations. For functional connectivity, lines connecting nodes indicate undirected functional connectivity, with blue depicting decreased connectivity. Weighted nodal degrees are represented in the heatmap tracks, depicting the cumulative sum of all its undirected connections. For effective connectivity, directed connections are displayed using arrowheads, with red lines indicating excitatory influence from one region to another in EC relative to controls, and blue lines representing inhibitory influence. Weighted nodal in-degree (incoming connections) and out-degree (outgoing connections) are also illustrated as heat maps.

***Across both counting and memory control conditions***. Notable findings include widespread lower functional connectivity throughout the brain, mostly between the brainstem and cortical networks, including the default-mode network (DMN), and between cortical networks, such as DMN-control network [CN], and CN-salience network (SN) during EC compared to both counting and memory control conditions.

***EC vs counting control condition.*** We observed reduced connectivity between the brainstem and somatomotor (SMN) and dorsal attention (DAN) networks (**Fig. 3a**) during EC compared to the counting condition.

***EC vs memory control condition.*** Interestingly, during EC, reduced functional connectivity between cortical networks such as the DAN, limbic network (LN), VN, and brainstem were notable (**Fig. 3b**). Additional reduced connectivity were found between the cerebellum and left subcortex.

### Effective connectivity: Directed network-based statistics (dNBS)

Results, including excitatory and inhibitory connections, and averaged weighted in-degree and out-degree are visualized in **Fig. 3c, d**. Here, we only provide a summary of the results.

***Across both counting and memory control conditions***. No similar excitatory or inhibitory connections were found across both control conditions compared to EC.

***EC vs counting control condition.*** No notable excitatory or inhibitory connections were found between EC and the counting control condition (**Fig. 3c**).

***EC vs memory control condition.*** Both excitatory and inhibitory connections were observed (**Fig. 3d**) during EC compared to the memory condition. Interestingly, inhibitory connections extend to nearly all brain networks, including the subcortex and brainstem. Specifically, we observed primarily excitatory connections to and from the left VN and brainstem, and inhibitory connections to and from the bilateral SMN, left SN, right CN and VN.

### Brain reorganization: Connectivity gradients

***Description and distribution of gradient values.*** The principal gradient depicted in the study aligns with previous research^30–34^, which identified a potentially hierarchical axis of functional connectivity similarity variance. This axis extends from unimodal regions, primarily located in the somatomotor cortex, to transmodal regions centered around the DMN and LN (**Fig. 4a**). Lower principal gradient values reflect greater functional connectivity similarity to unimodal cortex, whereas higher principal gradient values reflect greater functional connectivity similarity to transmodal cortex. We observed a wider range of gradient distribution across all networks. (**Fig. 4b**). Moreover, EC demonstrated an expansion of principal gradients compared to the two control conditions (**Fig. 4c**). Results for the bootstrapped t-tests at the ROI-level are presented in **Fig. 4d**, at the network-level **in Fig. 4e**. Significant results are also presented in the **Supplementary excel file**. Below, we summarize the results.

**Fig. 4.**
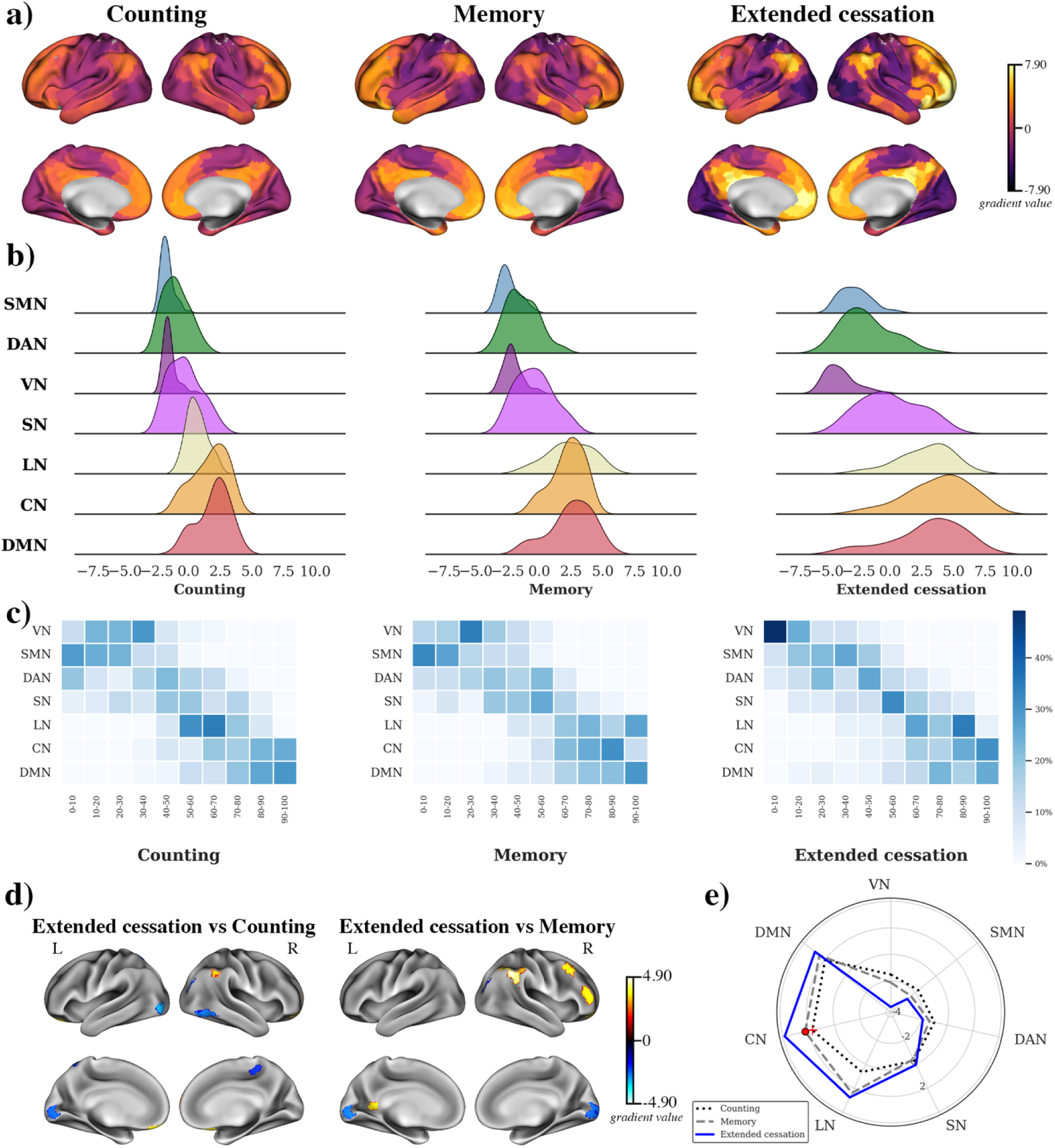
Differences in principal gradients between EC and control conditions. **(a)** Average gradients of each condition, demonstrating high gradient values in the transmodal regions during EC compared to control conditions. **(b)** Distribution of gradient values in each network during each condition. Expansion of the gradients were observed across all networks during EC. **(c)** Percentile of gradients along each network during each condition. Gradient values in the lowest 30 percentile during EC were distributed among unimodal regions (VN, SMN, DAN) while gradient values in the highest 30 percentile during EC were distributed among the LN, CN, and DMN. **(d)** Linear mixed models demonstrated that EC predominantly increases gradient values in the right parietal, and lower gradient values in the left visual cortex. **(e)** Linear mixed models demonstrated that EC exhibited higher gradient values in the CN. The red circle on the plot indicates significance between EC and the control condition. VN = visual network, SMN = somatomotor network, DAN = dorsal attention network, SN = salience network, LN = limbic network, CN = control network, DMN = default mode network.

***Across both counting and memory control conditions***. LMM at the network-level revealed that EC exhibited greater gradient values in the CN. At the ROI-level, EC demonstrated greater gradient values in the right frontal operculum insula, parietal cortex, and lateral PFC; and lower gradient values in the bilateral visual cortex within the visual network (VN). No additional significant gradient values were found.

### Cortical dynamics: Geometric eigenmodes

***Across both counting and memory control conditions***. LMM revealed no significant differences between EC and either counting or memory control conditions for mean, max, and total power or energy in any eigengroups. However, LMM revealed significant differences in main effects of task conditions between EC and control conditions for mean, max, or total power or energy (**Fig. 5**). Specifically, EC, compared to the counting condition, had lower max (b = - 0.09, p = 0.006) and total (b = -0.14, p < 0.001) power. Additionally, compared to the memory control condition, EC has lower mean (b = -0.15, p < 0.001), max (b = -0.31, p = 0.009), and total (b = -0.16, p < 0.001) energy, and lower mean (b = -0.32, p = 0.007), max (b = -0.21, p < 0.001), and total (b = -0.37, p = 0.001) power.

**Fig. 5.**
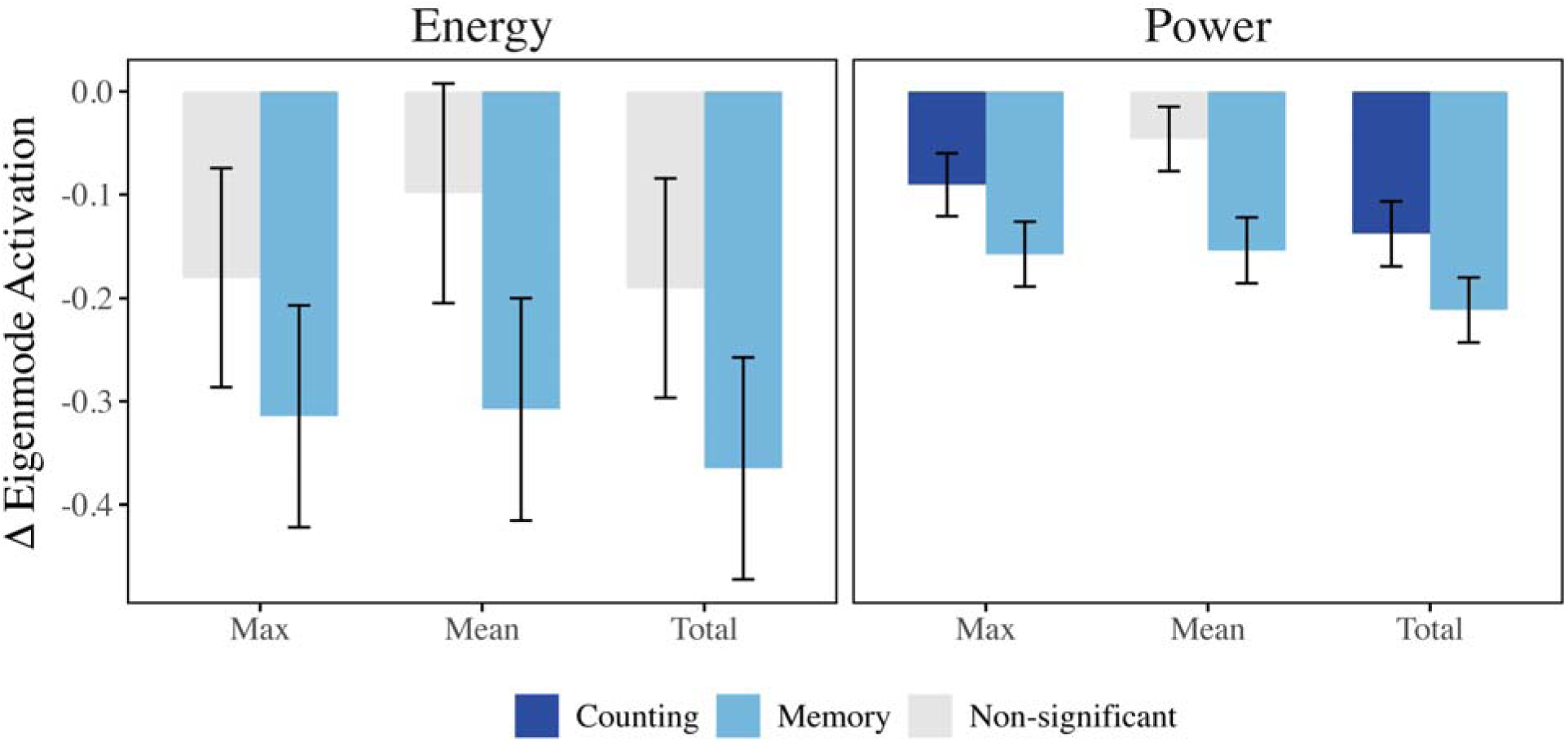
Differences in global eigenmode activations between EC and control conditions. Difference in global eigenmode activation for mean and max energy and power (averaged across eigenmodes), and total energy and power between EC - counting (dark blue) and EC - memory (cyan) conditions. Non-significant contrasts are colored grey. Error bars signify standard error.

### Neurotransmitter systems receptor density: Partial least squares correlation (PLSC)

PLSC extracted one significant latent variable relating neurotransmitter systems receptor densities to delta-weighted-degree maps, explaining 98.85% of the covariance between the two datasets (p_spin_ = 0.023, one-tailed). Additionally, a positive correlation was found between the latent variables (r = 0.42, p < 0.001, **Fig. 6a**). We also computed the loadings for each receptor (**Fig. 6b**) and delta-weighted-degree map (**Fig. 6c**), where positively loaded receptors co-vary with positively loaded delta-weighted-degree in positively scored brain regions and similarly for negative loadings^35^. Projecting the neurotransmitter systems receptor density patterns back onto the delta-weighted-degree weights reflects how well a brain area exhibits the receptor and delta-weighted-degree weighted pattern, referred to here as ‘receptor scores’ and ‘delta-weighted-degree scores’, respectively (**Fig. 6d, e**).

**Fig. 6.**
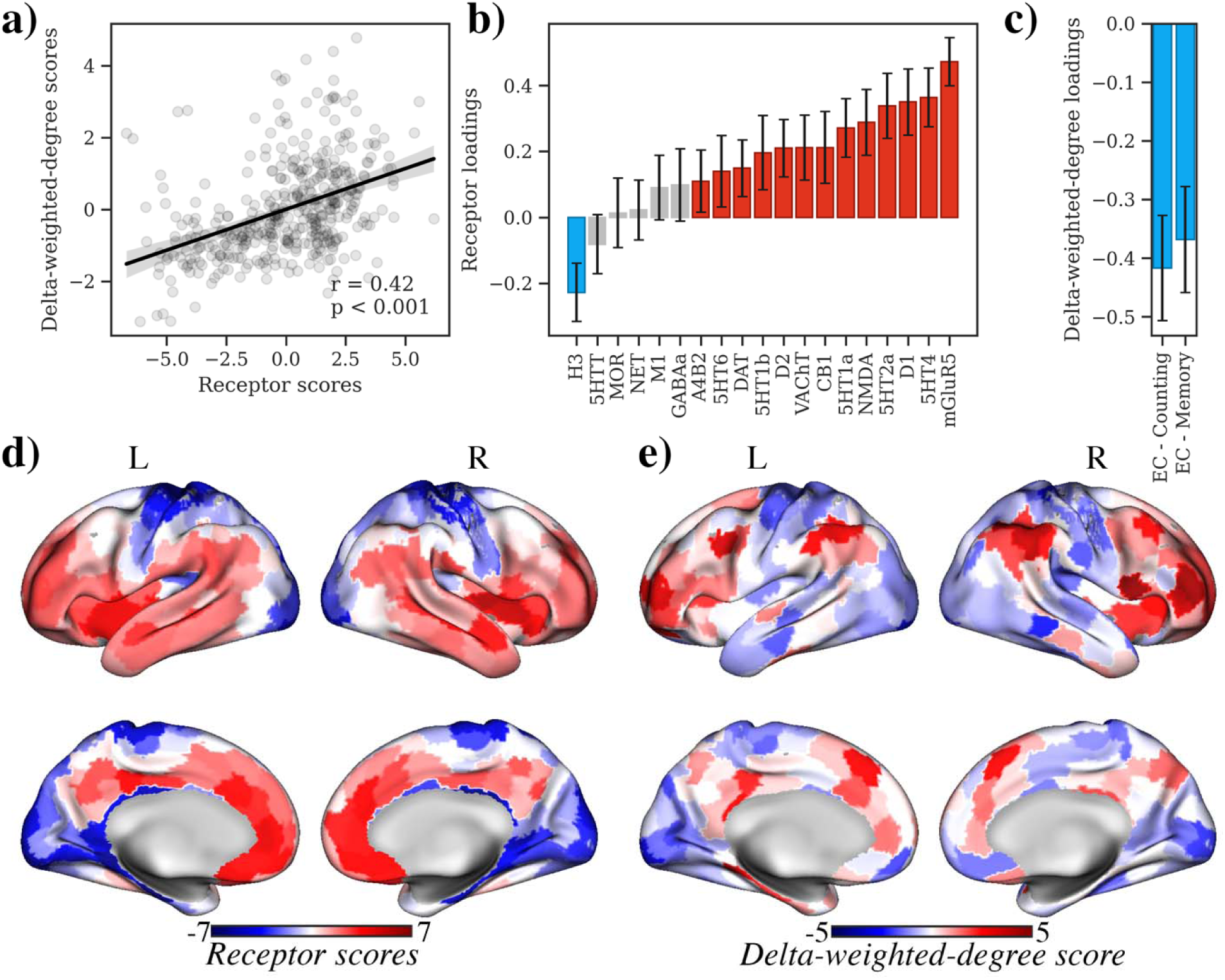
Spatial relationship between brain activity during EC and neurotransmitter receptor density. Only partial least squares (PLS) component 1 was significant. **(a)** Scatter plot shows the positive correlation between the latent receptor scores and latent delta-weighted-degree scores (r = 0.42, p < 0.001). **(b)** The bar chart displays the loadings of different neurotransmitter receptors on PLS component 1, with H receptors showing negative loading while mGluR_5_, 5HT4, D1, 5HT_2A_, and NMDA showing positive loadings. **(c)** The bar chart shows the loadings of delta-weighted-degree maps between task conditions, with negative loadings on the difference between EC and counting (EC - counting) and memory (EC - memory). **(d)** Spatial distribution of coactivation between receptors (‘receptor scores’) and (**e**) delta-weighted-degree (‘delta-weighted-degree scores’). Red areas indicate regions with positive delta-weighted-degree scores, while blue areas represent regions with negative delta-weighted-degree scores.

Metabotropic glutamate receptor 5 (mGluR_5_), serotonin 5HT_4_, and dopamine D_1_ displayed the greatest positive loadings while histamine H_3_ displayed the only negative loading (**Fig. 6b**). Non-significant loadings include serotonin transporter 5HTT, mu-opioid receptor MOR, norepinephrine transporter NET, and GABA_A_. Both EC-Counting and EC-Memory delta weighted-degree maps have similar negative loadings (**Fig. 6c**). This suggests that the combination of H_3_ receptor distributions co-vary with EC-related weighted-degree values in unimodal regions (**Fig. 6d**). EC reduces connectivity where mGluR /5HT /D receptors are abundant and increases connectivity where H receptors are abundant, relative to both control tasks. Examining the brain scores, transmodal regions (PFC, PCC, temporal areas), where mGluR /5HT /D /NMDA receptors are rich, decrease connectivity during EC relative to both controls (**Fig. 6e**). In contrast, primary sensory and motor cortices where H receptors are plenty, increase connectivity during EC. Collectively, these results demonstrate a direct link between whole-brain molecular receptor distributions and EC.

### Neurosynth data-driven cognitive terms decoding analysis

***Across both counting and memory control conditions***. Correlation analysis revealed that EC was positively associated with action, attention, fixation, gaze, planning, spatial attention, visual attention, and visual perception. These terms are associated with perception and planning. In contrast, negative associations were found with terms such as anxiety, arousal, eating, emotion, fear, loss, mood, pain, stress, and valence. These terms are associated with mental suffering and psychological distress. Results are visualized in **Fig. 2**.

***EC vs counting control condition.*** EC was additionally positively associated with imagery and object recognition.

***EC vs memory control condition.*** EC was additionally positively associated with selective attention and efficiency.

## Discussion

Using ultra-high-resolution 7T functional MRI, we present the first neural characterization of an extraordinary non-conscious state of extended cessation (EC). We used an intensive within-subject, dense-sampling design (N=3), analogous to the rigorous small-N primate study designs and small-N human case studies to prioritize internal validity and depth of individual-level analysis^36,37^.

Human consciousness is supported by integrated brain activity across functionally segregated areas^38^. Here we show that EC exhibits a distinctive reorganization of brain networks: higher brain activity in unimodal brain systems (VN, DAN), and temporal pol, lower activity in transmodal brain systems (DMN), as well as regions in the subcortex, brainstem, and cerebellum, expansion of the principal cortical gradient, enhancing segregation between unimodal and transmodal regions, and network–neurochemical coupling aligned with histaminergic H receptor topography. Furthermore, spatial meta-analytic decoding associates the EC pattern with cognitive processes associated with perception/attention and reduced mental suffering and psychological distress^39^. These results suggest a reconfiguration of the limbic network^40,41^, consistent with meditators’ post-EC reports and enduring psychological transformation of clarity, relief, and equanimity. These results offer novel insights into the neural correlates of EC, highlighting its potential to influence brain function as well as ultimately our understanding of consciousness and human flourishing, given the profound aftereffects reported.

Contrary to accounts that associate non-conscious states with global cortical suppression^42^ and gradient flattening and widespread integration loss^43^, EC showed higher local activity in unimodal sensory regions and reduced activity in transmodal regions and networks, specifically the DMN, and key subcortical and brainstem regions such as the caudate, thalamus, globus pallidus, and ventral tegmentum area. This reduced connectivity, notably in the basal ganglia^44–46^ and brainstem^47^ could be indicative of a indicating gating of access of sensation and percpetion^48^ and wakefulness^47^. These findings partially mimic patients having disorders of consciousness, such as those with minimally conscious state and unresponsive wakefulness syndrome^49^, who also demonstrated reduced brain activity in the DMN. In contrast, the principal gradient expands (unimodal values become more negative; transmodal values become more positive) and the distribution was flatter across all networks in EC, suggesting specialization of brain functions during EC. Unlike non-conscious states due to pharmacological interventions or disorders of consciousness, EC is endogenous, volitional, and followed by equanimity rather than disorientation. These contrasts support the view that multiple routes to non-conscious states exist, such as pharmacologic/global suppression versus selective decoupling, with divergent network consequences.

Two most prominent theories of consciousness are GNW and IIT. GNW posits that consciousness depends on long-range recurrent broadcast across frontoparietal hubs and non-consciousness accompanies reduced feedback and integration (e.g., anesthesia)^27^. EC is partly consistent with this view wherein DMN and thalamic suppression imply diminished recurrent broadcast to the workspace but constrains GNW because unimodal systems (VN/DAN) remain engaged and the principal gradient polarizes rather than collapses. Thus, selective decoupling of recurrent access to transmodal hubs, rather than frontoparietal shutdown, can suffice to suspend awareness. In contrast, IIT predicts that non-conscious states reflects reduced integration/differentiation, which often manifests empirically as flattened principal gradients and loss of complex spatial modes in pharmacological interventions and disorders of consciousness^28,43,50^. EC shows the opposite pattern: gradient expansion, stronger unimodal– transmodal segregation in the absence of awareness, and reduced lower-order brain modes in EC. This directly challenges IIT, indicating that excessive segregation rather than loss off integration can also yield non-conscious states. Finally, according to AIF, loss of consciousness can arise from down-weighting the precision of sensory prediction errors^29^. Surprisingly, compared to GNW and IIT, EC supports AIF on multiple fronts. The reduced activity in the thalamus and brainstem together with increased activity in unimodal systems is consistent with precision gating. Additionally, network–neurochemical coupling to histamine H receptor topography in unimodal cortex could explain neuromodulation on precision control^35^. Finlly, post-EC sensory clarity and equanimity is consistent with a transient precision overshoot, a brief increase in the gain assigned to sensory prediction errors which yields unusually vivid perception while self-referential priors remain down-weighted. In short, EC supports precision-gating (AIF), refines GNW by dissociating recurrent access from generic frontoparietal activity, and challenges IIT by demonstrating non-conscious states with increased hierarchical segregation rather than uniform flattening.

A unique contribution of this study is the link between EC and neurotransmitter receptor topography. Brain activity during EC co-varies with H receptors receptor topography. As presynaptic autoreceptors and heteroreceptors, H receptors regulate histamine release and suppress acetylcholine, dopamine, and norepinephrine, shaping arousal, attention, and sensory filtering^51–53^. EC’s coupling to H -dense unimodal cortices and reduced engagement of transmodal hubs rich in mGluR /NMDA/D may allow habitual cognitive constraints to relax, enabling underlying intrinsic brain dynamics to emerge^54^ and modulate the type of conscious experience^55^. This shift likely allows practitioners to re-engage perception with heightened equanimity and reduced cognitive bias, providing optimal conditions for deep meditative insight. Notably, similar decreases in transmodal network activity have been observed during ACAM^2^ and AIIM^56^. Upon exiting EC, unimodal networks enable a rapid, equanimous re-engagement of sensory processes, giving rise to the vivid clarity, equanimity, and deep insight into the phenomenology of the mind practitioners describe.

EC may be best understood as a “reset” of consciousness, in which self-referential loops are silenced, perceptual systems reorganized, and affective load attenuated. Thus, beyond the field of cognitive neuroscience, EC offers a radical empirical model for advancing mental health, wellbeing, and human flourishing. The transformative aftereffects of EC—including deep sense of relief, clarity, equanimity, and reductions in repetitive thought, self-talk, and craving—map onto diminished self-referential processing and enhanced perceptual networks, according to the decoded meta-analytic cognitive maps from Neurosynth. These neural and experiential changes directly address components of psychological suffering^10^. Finally, the neural signatures of EC could serve as biomarkers for advanced meditation outcomes, guiding machine-learning-based personalization of practice^57^ or neuromodulation protocols to target specific meditation outcomes, such as kindness, concentration, or acceptance^58^. Thus, EC point towards novel strategies for facilitating increasingly effective mental health interventions^59^.

This study has considerable strengths including a whole-brain approach at high spatial resolution using ultra-high-resolution fMRI data with stringent methodological approaches. Nevertheless, there are several important limitations to consider. First, as this study is the first to investigate extended cessation, specifically EC, our results should be considered an initial step that should be replicated. While we compared EC to non-meditative states, a natural extension would be to compare EC to other meditative states to elucidate similarities and differences between EC and advanced forms of mediation. The absence of consciousness and self-reports during EC poses a challenge in understanding the subjective non-experience and its correlation with the neuroimaging findings. However, Neurosynth decoding offered robust evidence linking EC brain state to enhanced perceptual clarity, sensitivity, and the absence of mental suffering. These findings strongly suggest that significant brain reorganization occurs during EC to a degree that supports these subjective subsequent phenomena.

In conclusion, our results provide the first evidence for the neural correlates of extended cessation. Extended cessation is marked by higher activity in unimodal regions with downregulation of transmodal and thalamic regions, expansion of cortical gradients, and reduced low-order global eigenmodes. Our results supports AIF, constraints GNW, and challenges IIT. These findings not only deepen our understanding of extended cessation but also illuminate fundamental mechanisms underlying human consciousness. More broadly, EC reveals a selective “reset” capacity of the mind that attenuates self-referential burden while sharpening perceptual engagement, opening tractable neuromodulatory and behavioral routes to reduce suffering and advance human flourishing.

## Materials and Methods

### Participants

Participants were three male advanced meditators, Subject 2 (Male, age 32 at time of data collection), Subject 3 (age 46 at time of data collection), and Subject 4 (age 66 at time of data collection). Important to the current study, the participants are advanced meditators and reported being able to incline towards EC as the target of meditation. Subject 2 had over 19 years of meditation experience with total meditation experience of an estimated at least 15,000 hours. Subject 3 had over 4 years of meditation experience, and an estimated total practice amount at least of 9,000 hours. Subject 4 had over 50 years of meditation experience with total meditation experience of at least 35,000 hours. Lifetime practice hours are approximate self-reports based on duration and frequency of weekly practice and retreats. We also note that lifetime hours are not necessarily a direct measure of meditative expertise. The Mass General Brigham IRB approved the study, and the participants provided informed consent.

### Experimental design

#### Extended cessation (EC)

FMRI data during EC was collected as part of a larger study of advanced meditation practices. In this advanced practice of EC, participants have trained extensively to develop the precision to enter and exit EC at specific, predefined times. This level of mastery reflects not only the ability to achieve a deep cessation state but also control over the duration and timing of the state. The participants performed EC meditation as the last meditation session after all other MRI sessions (structural scans, advanced concentrative absorption meditation, advanced insight meditation, non-meditative control conditions) were completed. The total amount of fMRI data collected for EC for each participant was approximately 45 minutes across three sessions. We asked the participant to meditate, enter, and maintain EC for 15 minutes during each session.

An EC run started with the participant making an intention to stay in EC for only 15 minutes. Then the participant started meditating at the start of the EC run. As some participants indicated that it was not possible for them to self-report entrance to this state, we did not attempt to mark when the participant entered EC. The run ended when the participant exited out of EC and indicated the exit with a button press. Retrospectively, we asked participants to estimate the duration it took them to enter EC if they did not indicate with a button press initially. After completing the meditation task, all participants completed a standardized EC phenomenology questionnaire—designed to reflect how they typically practice and experience EC—covering preparation, onset, absence of consciousness, exit, and afterglow. Two participants provided their phenomenology responses after data collection had formally concluded, and their delayed reports have been included in the dataset.

#### Non-meditative control conditions

We developed two non-meditative control conditions (hereon control conditions) that were used to compare against EC. These control conditions were designed to engage the participant’s mind with non-meditative mental activities. These conditions were carefully selected to be sufficiently engaging to not induce meditative states. We did not use a resting-state control condition, since experienced meditators may enter meditative states during this period^60^.

The two different control conditions implemented in this study were: (1) a memory control condition in which participants were asked to reminisce the events of the past two weeks and narrate them sub-vocally in their minds, closing their eyes and without moving their lips, for 8 minutes (min); and a (2) counting control condition, where the participants were asked to mentally count down in decrements of 5 from 10,000 for 8 min, closing their eyes and without moving their lips. We collected two runs for each control condition, which provided 16 mins data for each control condition. However, for participant ABC, only a single run for each control condition as part of another study.

### Neuroimaging acquisition

Neuroimaging was acquired using a 7T MR scanner (SIEMENS MAGNETOM Terra) using a 32-channel head coil. Functional imaging was performed using a single-shot two-dimensional echo planar imaging sequence with T2*-weighted BOLD-sensitive MRI, repetition time (TR) = 2.9 sec, echo time (TE) = 30 ms, flip angle (FA) = 75°, field of view (FOV) = [189 x 255], matrix = [172 x 232], GRAPPA factor = 3, voxel size = 1.1 x 1.1 x 1.1 mm^3^, 126 slices, interslice distance = 0 mm, bandwidth = 1540 Hz/px, echo spacing = 0.75 ms. Slice acquisitions were acquired for the whole brain, with interleaved slices, sagittal orientation, and anterior-to-posterior phase encoding. Opposite phase-encoded (i.e., posterior-to-anterior) slices with the same parameters were also acquired to perform distortion correction.

Whole-brain T1-weighted structural images were acquired as follows: TR = 2.53 sec, TE = 1.65 ms, inversion time = 1.1 sec, flip angle = 7°, 0.8mm isotropic resolution, FOV = 240 x 240, GRAPPA factor = 2, bandwidth = 1200 Hz/Px. The participant’s physiological (i.e., heart rate using pulse oximetry and respiration using breathing bellows) signal recordings were collected throughout the scanning session.

Each neuroimaging session started each time the participant’s position was localized in the scanner and lasted until the next localization was conducted, usually after the participant left the scanner to rest, typically after two hours.

### Neuroimaging preprocessing

Preprocessing steps were conducted at the level of MRI session and for all runs within that session, similar to our previous protocol, using AFNI^2^. Details of the preprocessing steps are described in our previous study^2^. In short, steps consisted of (1) de-spiking; (2) RETROspective Image CORrection (RETROICOR)^61^ to regress out the effects of physiological (cardiac and respiratory) noise on data; (3) slice time correction; (4) distortion correction using opposite phase-encoded EPI; (5) motion correction; and (6) registering the anatomical dataset (T1) to a standard (MNI152_2009) template. Additional preprocessing steps included: (7) scrubbing any volume with motion > 0.3mm and had more than 5% outlier voxels; and (8) regressing out eroded cerebrospinal fluid (CSF) mask time course and motion parameters (3 translations, 3 rotations) per run, and band-pass filtering (0.01 – 0.1Hz). Each fMRI run was then segmented into one-minute segments for statistical analyses, similar to our previous work on advanced meditation ^2^.

### Regions of interest (ROI)

We used four different parcellation/segmentation schemes to define ROIs for subsequent whole-brain analyses: (1) Schaefer-400 parcellation atlas for cortical areas^62^; (2) 34-region Tian subcortex atlas for subcortical regions^63^; (3) 54 Bianciardi brainstem atlas^64^; and (4) 10 Multi-Domain Task Battery (MDTB) functional cerebellar atlas^65^. In total our parcellation/segmentation yielded 498 ROIs across the brain. For functional and effective connectivity analyses specifically, we opted to use the Schaefer-100 parcellation atlas for cortical areas instead of the 400 parcellation scheme^62^ and the 16-region Tian subcortex atlas for subcortical regions^63^. This decision was made to reduce the number of ROIs, thereby facilitating the interpretation of whole-brain effective connectivity patterns and enabling direct comparability between functional and effective connectivity results, while still maintaining a reasonably high spatial resolution for cortical regions.

### Neuroimaging analyses

#### Regional homogeneity analysis

Regional homogeneity (ReHo) is a measure of similarity in temporal activation pattern of a voxel and its nearby voxels^66^. This measure of local functional connectivity within brain regions is a close derivative of underlying brain activity^66^. A higher ReHo value indicates stronger synchronization of local brain regions, indicating greater functional connectivity in that region and thus an index of greater brain activity. In this study, we defined a cluster size of 27 voxels and calculated the ReHo values for each fMRI segment. We standardized the ReHo values before smoothing the standardized ReHo maps using a 2 mm full-width half-maximum (FWHM) kernel. The standardized ReHo values were then parcellated using the four different parcellation/segmentation schemes^62–65^, yielding 498 ReHo values for each fMRI segment.

#### Functional connectivity

Data was smoothed using a 2 mm FWHM as we were interested in connectivity of smaller brain regions such as the brainstem. Then for each segment, whole-brain time series of the 180 brain regions as defined previously was extracted, followed by computation of 180x180 functional connectivity matrices using Pearson’s correlation.

#### Effective connectivity: Regression dynamic causal modeling (rDCM)

Whole-brain effective connectivity was assessed using regression dynamic causal modeling (rDCM). rDCM, a variant of DCM for fMRI, enables effective connectivity analyses across whole-brain networks with computational efficiency^67–71^ without sacrificing reliability^72^. The validity of this procedure has been established in both task-based and resting-state fMRI studies^67,68,70,72,73^. Specifically, rDCM incorporates modifications to DCM^72^ including: (1) translation of equations into the frequency domain using Fourier transformation; (2) replacement of nonlinear hemodynamic models with linear functions; (3) mean-field approximation across regions; and (4) specification of priors for neuronal parameters and noise precision. These modifications transform linear DCM into Bayesian linear regression in the frequency domain, which ultimately facilitates highly efficient DCM model inference^70,73^.

To compute effective connectivity, preprocessed data at step 6 (normalization) was first smoothed using a 2 mm FWHM. Time series of each ROI were then extracted after regressing out 6 motion parameters, white matter and CSF signals, with discrete cosine basis set with frequency characteristics of resting-state brain dynamics (0.01– 0.1 Hz). Whole-brain effective connectivity was then calculated using rDCM for resting-state fMRI^68^ to examine whole-brain effective connectivity^69,74–77^. First-level rDCM models for 180 ROIs were then computed using the rDCM module within the Translational Algorithms for Psychiatry Advancing Science (TAPAS) toolbox^78^.

#### Principal gradient-mapping

To map the gradients, we first computed a template gradient through a multi-step process. First, data was smoothed using a 6 mm FWHM kernel, followed by computation of functional connectivity matrices for each segment using the Schaefer 400-regions 7-network atlas using Pearson’s correlation, thus producing a 400×400 functional connectivity matrix. These individual matrices were then averaged across all segments, resulting in a mean functional connectivity matrix. Subsequently, gradients for this mean matrix were derived by thresholding the mean correlation matrix at 90% sparsity to retain only the strongest connections, followed by generating a similarity matrix using cosine similarity to capture the connectivity pattern similarities among ROIs. The resulting similarity matrix served as input for the diffusion map embedding algorithm, from which the principal gradients were extracted. We computed a template gradient from an out-of-sample dataset of 134 subjects from the HCP dataset (the validation cohort used by^79^).

Following template gradient computation, alignment with individual segment gradients was performed. The principal gradient values for each fMRI segment were computed based on their respective functional connectivity matrices and then aligned with the template gradient using Procrustes rotations with 10 iterations to ensure optimal comparison across datasets. All analyses described herein were conducted using the BrainSpace toolbox implemented in Python^79^. Using the gradient values obtained from gradient alignment, we computed the average network gradient values for each network by averaging the gradient values of all the parcels within that network.

#### Cortical dynamics: Geometric eigenmodes

To decompose fMRI activity into 200 frequency-specific eigenmodes, we used geometric eigenmodes derived from a population-averaged midsurface thickness template of the neocortical surface from Pang et al., wherein the methods to generate these geometric eigenmode templates are also described^80^. Briefly, the geometric eigenmodes were computed by constructing the Laplace-Beltrami operator from the cortical mesh and solving the eigenvalue problem. The resulting eigenvalues are ordered according to spatial frequency/wavelength of each mode, where mode 1 has the longest wavelength (lowest frequency) while higher modes have shorter wavelengths (higher frequency). Conversely, eigenmodes are the complete basis set—a weighted sum of varying-wavelength modes. Lower-numbered modes capture large-scale spatial patterns (e.g. left/right hemisphere, anterior/posterior regions) while higher-number modes correspond to smaller-scale spatial patterns where functional properties of nearby regions vary independent of proximity.

Using the geometric eigenmodes, we decomposed functional MRI data to reveal emerging cortical spatiotemporal dynamic patterns by examining the contribution of each eigenmode to the cortical activity at each time instance. At each time point of each fMRI time courses, fMRI data were projected onto each of the 200 geometric eigenmodes, yielding the temporal activity of the particular eigenmode in a method described previously^81,82^. Subsequently, the temporal activation of eigenmodes were analyzed in terms of their power (the strength of an eigenmode’s activation at a given fMRI time instance: |ω_k_(t_i_)|) and energy (the eigenmode’s frequency-weighted contribution, estimated by weighting the square of the eigenmode’s strength of activation by the square of its corresponding eigenvalue (λ*_k_*) at each time point: |ω_k_(t_i_)|^2^ λ*_k_*^2^) at each fMRI time point. Mean power and energy were calculated by averaging the power and energy of each eigenmode across all time points in a single segment (i.e. yielding 200 mean power and energy values for 200 modes per fMRI segment). Maximum power and energy were calculated as the maximum spectra value for a given eigenmode across all time points within a segment (i.e. yielding 200 maximum power and energy values for 200 modes per fMRI segment). Unless otherwise noted, all reported maximum and mean power and energy values are reported on a log-scale^82^. Brain total power and total energy were computed by summing power and energy across all eigenmodes for each time point and then averaging all time points within a segment.

#### Influence of neurotransmitter systems receptor density: Positron Emission Tomography (PET) neurotransmitter systems receptor density maps

We used a recently made available public neurotransmitter systems receptor density maps for 19 receptors and transporters, across 9 neurotransmitter systems by Hansen and colleagues at https://github.com/netneurolab/hansen_receptors^35^. These include dopamine (D_1_, D_2_, DAT) noradrenaline (NAT), serotonin(5-HT_1A_, 5-HT_1B_, 5-HT_2A_, 5-HT_4_, 5-HT_6_, 5-HTT), acetylcholine (α_4_β_2_, M_1_, VAChT), glutamate (mGluR_5_, NMDA), GABA (GABA_A_), histamine (H_3_), cannabinoid (CB_1_), and opioid (MOR) receptors. Details regarding the neurotransmitter systems receptor density maps, each PET dataset, and their respective acquisition and limitations can be found in ref^35^. Volumetric PET images were registered to the MNI-ICBM 152 nonlinear 2009 (version c, asymmetric) template, averaged across participants within each study, and then parcellated and receptors/transporters with more than one mean image of the same tracer were combined using a weighted average^35^.

#### Decoding of brain activity using data-driven Neurosynth cognitive maps

To provide a complementary approach for analyzing brain activity patterns with behavior in the absence of behavioral data, we examined the spatial similarity of ReHo maps with 123 meta-analytic brain maps derived from the Neurosynth database. Neurosynth generates probabilistic measurements that can be interpreted as a quantitative representation of the association between regional fluctuations in activity and psychological processes. Although Neurosynth’s database contains over 1,000 terms, we narrowed our focus to terms related to cognition and behavior, drawing from previous research^35,83,84^. Our selected set of 123 terms encompasses broad categories, ranging from attention and emotion, to more specific processes such as visual attention and episodic memory. The terms also include basic behaviors (e.g., eating and sleep) and emotional states (e.g., fear and anxiety).

### Statistical analysis

#### ReHo brain activity

Linear mixed-effects models were used to analyze ReHo differences between EC and control conditions. The model included task condition as a fixed effect and random intercepts for participant and for segments nested within subject. Corrections for multiple comparisons across models were applied using false-discovery rate (FDR) across all 498 ROIs. We used Dunnett’s test to compare the difference between EC and control tasks, with EC as the reference group controlling for multiple comparisons using Bonferroni correction for k = 2 comparisons. For ease of interpretation, all coefficients were flipped to retrieve the contrast of EC against the control conditions. Analyses were conducted in *R*^85^ loading on R Studio v2022.12.0.353.20^86^.

#### Functional connectivity: Network-based statistics (NBS)

NBS was used to explore functional connectivity differences between EC and control conditions in the whole-brain network containing 198 nodes and 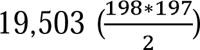 unique edges^87^. NBS is a nonparametric statistical method used to perform mass-univariate statistical tests for multiple graph comparisons. NBS identifies interconnected subnetworks exhibiting significant differences in functional connectivity between groups or conditions. Instead of treating connections independently, NBS leverages the inherent interconnectedness of the brain by identifying clusters of connections collectively differing across groups. This enables sensitive detection of subtle yet interconnected alterations in whole-brain functional connectivity. T-scores were computed for each pairwise connection under contrasts EC > control and EC < control separately, with additional separate analyses for counting and memory control conditions. Each test was conducted at the primary threshold (t = 3.1, p = 0.025) to select the suprathreshold connections for which the t-scores exceeded the primary t-value. We used p = 0.025 to accommodate two contrasts per control comparison (EC > control and EC < control). Connected components where the EC showed higher or lower functional connectivity than the control conditions were identified, and the number of the edges or their size was stored. A permutation test was then performed to determine the significance of each component by comparing its size to the distribution of maximum component sizes under random group assignments (5000 permutations). Corrected p-values were computed as the proportion of permutations with larger maximum components than the observed size. NBS enabled sensitive detection of interconnected functional connectivity alterations in EC by leveraging the inherent network structure of the brain and rigorously accounting for multiple comparisons.

#### Effective connectivity: Directed network-based statistics (dNBS)

dNBS, an extension of NBS, was used to analyze effective connectivity differences between EC and control conditions. We investigated two contrast, EC > control and EC < control separately, with additional separate analyses for counting and memory control conditions. The statistical significance of the outcomes was determined using a primary threshold of t = 3.1 (p = 0.025) and 5000 permutations, following the same procedures as in the conventional NBS approach. The key distinction between dNBS and NBS lies in the fact that dNBS analyzes twice the number of edges compared to NBS, accommodating the directional nature of effective connectivity measures.

#### Brain organization: Connectivity gradients

Linear mixed-effects models were used to analyze cortical principal gradient differences between EC and control conditions. These tests were performed at both the network-level, encompassing 7 cortical networks, and at the ROI-level, comprising 400 ROIs. This dual approach allowed for a robust examination of gradient variations across different scales of cortical organization. Corrections for multiple comparisons were applied using false-discovery rate across the 7 networks and 400 ROIs respectively.

#### Cortical dynamics of EC: Geometric eigenmode power and energy

To analyze differences in frequency-specific eigenmode mean power and energy between EC and control conditions while minimizing multiple comparisons, we utilized 15 eigengroups: grouping of eigenmodes according to spatial frequency^80,88^. Eigengroups provide a way to aggregate degenerate solutions from the solving of the eigendecomposition such that eigenmodes with the same number of nodal lines and wavelengths are grouped together. This method allows for investigation of eigenmode metrics that are less affected by noise^89,90^. Using this approach, each eigengroup’s power and energy were then compared as reference values to both counting and memory control conditions using linear mixed-effects models. The resulting *p*-values were FDR corrected across both control condition comparisons. Finally, linear regression was used to examine mean power and energy differences between EC and control conditions.

#### Influence of neurotransmitter systems receptor density: Partial least squares correlation

We used partial least squares correlation (PLSC) analysis to explore the multivariate relationship between neurotransmitter systems receptor densities and EC delta-weighted-degree connectivity maps. PLSC is an unsupervised multivariate statistical technique that decomposes two datasets into orthogonal sets of latent variables with the goal of maximizing covariance between them^91^. In this context, the latent variables consist of receptor weights, EC-control weighted-degree weights and a singular value that represents the covariance between receptor distributions and weighted-degree connectivity that is explained by the latent variable.

EC delta-weighted-degree maps were created by first thresholding all negative-weighted connectivity and self-connections to zero, followed by summing each region’s positive connection weights to all other regions. we only use cortical atlas for this analysis, similar to previous research on chemoarchitecture of the brain^35,83^. Segment-level weighted-degree vectors were then averaged across all segments within each task, producing a single weighted-degree profile per participant per task. For each participant, the delta-weighted-degree map was calculated as the difference between EC and each control task, i.e., EC-Counting, and EC-memory. Finally, participant-level delta-weighted-degree profiles were then averaged across the three participants to obtain the two delta-weighted-degree maps, EC-Counting, and EC-memory.

By projecting the original receptor density and weighted-degree data onto the corresponding latent variable weights, we computed receptor and EC-difference scores, assigning each brain region a specific score for both. Receptor loadings were then calculated as the Pearson correlation between neurotransmitter systems receptor densities and receptor scores, with a similar process used to compute EC-difference loadings (weighted-degree map and EC-difference scores). Importantly, PLSC does not (1) infer causal links between receptors and EC, (2) establish univariate associations between specific receptors and EC, or (3) rule out the possibility of additional connections between receptors and EC.

We assessed the significance of each latent variable’s singular value using permutation spin-testing^92–94^. For each map, parcel coordinates were projected onto the spherical surface and then randomly rotated and original parcels were reassigned the value of the closest rotated parcel (10,000 repetitions)^95^. In addition to preserving the distribution of cortical values, this null model also preserves the spatial autocorrelation present in the data.

#### Contextualization of brain activity with Neurosynth cognitive maps

We used the *continuous.CorrelationDecoder* function from the NiMARE python package to contextualization of brain activity with Neurosynth cognitive maps^96^. We first fetched Neurosynth dataset and converted it to NiMARE dataset. Following, we retrieved the 123 cognitive terms from the database. We also computed delta-ReHo maps (EC-counting, and EC-memory) for decoding. Finally, we used the *CorrelationDecoder* function to decode the association between the unthresholded delta-ReHo maps and cognitive maps.

## Supporting information

Supplementary dictionary

## Author contributions

Conceptualization: MDS; Funding Acquisition: MDS; Methodology: MDS, TS; Data Collection: MDS, TS, RP, WFZY; Software: WFZY, GM, IB, RP; Formal Analysis: WFZY, AK, KA, GM, IB; Writing – Original Draft: WFZY and MDS; Writing – Review & Editing: WFZY, MDS, AK, KA, RP, and TS.

## Funding information

Dr. Sacchet and the Meditation Research Program are supported by the Dimension Giving Fund, Tan Teo Charitable Foundation, and additional individual donors. Dr. Abellaneda-Pérez was financially supported by a Juan de la Cierva research grant (FJC2021-047380-I) of the Spanish Ministry of Science and Innovation. Dr. Terje Sparby is supported by Software AG Stiftung.

## Competing interests

All authors declare no competing interests.

## Data and materials availability

Data may be requested from the corresponding author and is subject to the Massachusetts General Hospital Institutional Review Board’s guidelines and approval. All code used in this study may be made available by request from the corresponding author.

## Notes

### Competing Interest Statement

The authors have declared no competing interest.

### Summary of Updates

Corrected minor typos in the manuscript.

