## Supplementary dictionary for "Endogenous suspension and reset of consciousness: 7T fMRI brain mapping of the extended cessation meditative endpoint"

This dictionary file provides information regarding the contents within each Excel tab, summarizing significant findings presented in the paper.

**ReHo Tab:** Contains significant Regional Homogeneity (ReHo) analysis results related to extended cessation across counting and memory tasks. Included are Linear Mixed Model (LMM) statistics such as chi-square values, parameter estimates, standard errors, t-values, adjusted p-values with Bonferroni correction for k=2 comparisons, and FDR-adjusted chi-square p-values across 498 Regios of Interest (ROIs).

- **Control Condition:** Experimental condition used as the control (i.e., EC – memory and EC - counting).
- **Atlas:** Brain atlas used to define the ROI (Schaefer, Tian, Bianciardi, MDTB).
- **ROI:** Specific Region of Interest under analysis.
- **Network:** Functional brain network assignment of the ROI (e.g., DMN, FPN).
- **singular:** Singular value used in model fitting (likely for regularization or dimensionality).
- **chisq:** Chi-square test statistic from the linear mixed model.
- **chisq.pval:** Raw p-value for the chi-square test.
- **Estimate:** Estimated effect size from the model.
- **SE:** Standard Error of the estimate.
- **TVAL:** T-statistic value from the model.
- **PVAL_ADJUSTED:** Bonferroni-adjusted p-value for t-test comparison (k=2).
- **CHISQ_PVAL_ADJUSTED:** FDR-adjusted p-value for chi-square test (corrected across 498 ROI tests).

**FC EC-Counting and FC EC-Memory Tabs:** Present significant functional connectivity findings between 180 ROIs, derived from Network-Based Statistics (NBS) analyses specific to EC - counting and EC - memory.

- **ROI:** Seed region for the connectivity matrix.
- **All other columns:** Target ROIs.
- **Values:** T-statistic of pairwise functional connectivity.

**rDCM EC-Memory Tab:** Provides significant effective connectivity outcomes between 180 ROIs identified via Directed Network-Based Statistics analyses using regression Dynamic Causal Modeling (rDCM) specifically for memory tasks. ***Note that no significant findings were found for EC-counting.***

- **ROI**: Source ROI for effective connectivity.
- **Remaining columns:** Target ROIs.
- **Values:** T-statistic.

**Gradients - Networks Tab:** Displays significant network-level findings from Principal Gradient analyses conducted through Linear Mixed Models.

- **Control Condition:** Experimental condition used as the control (i.e., EC – memory and EC - counting).
- **Network:** Brain network label (e.g., DMN).
- **Remaining columns:** Same LMM statistics as in ReHo (e.g., chisq, Estimate, PVAL_ADJUSTED).

**Gradients - ROIs Tab:** Lists significant ROI-level findings from Principal Gradient analyses, consistent with the ReHo tab details.

- Same columns as above but at ROI-level.
- **ROI:** Specific brain region.
- **Network:** Assigned network of the ROI.

**Eigenmodes - Regression Tab:** Documents significant metrics including maximum, total, mean power, and energy from the Geometric Eigenmodes analyses.

- **Control Condition:** Group/task comparison.
- **Effect:** Name of experimental effect.
- **Metric:** Power/energy measure.
- **Stat:** Statistic computed for the metric (e.g., Max power, Total power, Mean power).
- **Remaining columns:** LMM statistics (Same as the ReHo tab).

**Neurosynth Tab:** Features comprehensive correlations between ReHo values and 123 cognitive neuroscience terms from the Neurosynth database.

- **Feature:** Cognitive term from the Neurosynth meta-analytic database (e.g., "attention", "memory").
- **EC-Counting r / EC-Memory r:** Pearson's correlation (r) between ReHo and Neurosynth feature map during for EC-counting and EC-memory contrasts.

**PLSC Scores Tab:** Presents delta-degree connectivity scores and receptor scores derived from Partial Least Squares Correlation (PLSC) analyses.

- **ROI:** Brain region.
- **Network:** Functional network label.
- **Hemisphere:** LH (left), RH (right).
- **LV1-X-scores / LV2-X-scores:** Delta-degree-connectivity scores.
- **LV1-Y-scores / LV2-Y-scores:** receptor profile scores.
